# High efficient chromatin conformation capture without pre-enrichment (HiChew) in single cells

**DOI:** 10.1101/2024.06.25.600609

**Authors:** Zhichao Chen, Yeming Xie, Chen Tan, Fengying Ruan, Meng Luo, Chen Zhang, Mei Guo, Yitong Fang, Chong Tang

## Abstract

This study presents HiChew, a cutting-edge technique for high-efficiency chromatin conformation capture in single cells, without the need for pre-enrichment. This unique approach minimizes the risk of cell or DNA loss. When compared to Dip-C, HiChew captures valid pairs with 4-8 times more efficiency, reducing wastage and saving significant sequencing budget. Furthermore, HiChew delivers a lower false positive ratio, ensuring data accuracy. It also achieves more contacts per cell, enhancing resolution in single cell. HiChew’s superior performance not only enhances single-cell Hi-C but also streamlines conventional Hi-C, making it more robust than conventional HiC methods. This study also unveils a fascinating mechanism of gene activation in the B compartment of chromatin, providing insight into the elusive aspect of gene expression within this region.

## Introduction

Chromatin conformation capture methods are useful tools for exploring the three-dimensional structure of chromatin in cells. These methods enable researchers to observe how the genome is arranged in space, which is essential for comprehending gene expression and regulation.

Chromatin conformation capture^1^ is a technique that involves crosslinking genomic DNA, digesting it with a restriction enzyme, end repair to incorporate the biotin, and then ligating the resulting fragments. To enrich the ligated products, we usually adopt the streptavidin enrichment method, which helps to generate more valid pairs. In conventional Hi-C, the ligation occurs between proximal fragments, providing information about the spatial proximity of different regions of the genome. Finally, we sequence the resulting library to identify the interactions between different regions of the genome.

However, because the genome structure can vary greatly between different cells, the conformation observed is actually an average of what is seen in various cells. In order to observe the unique structures of individual cells and avoid any interference from neighboring cells, scientists have developed single-cell Hi-C^2^. This method offers the advantage of a more precise observation of the dynamic nature of chromatin structures. There are two main styles of methods in the single-cell Hi-C field: enrichment methods and non-enrichment methods.

In the initial stages, scientists developed the first-generation single-cell Hi-C protocol, which was similar to conventional Hi-C but optimized for minimal input DNA^2^. However, biotin enrichment resulted in significant DNA loss, most likely due to inefficient biotin incorporation or loss of biotin binding. Subsequently, scientists developed the biotin incorporation-free method, ChIA-PET^3^, which utilized a biotin linker to ligate between the two neighboring fragments (in junction position), eliminating the inefficient biotin incorporation. However, the adapter required to ligate both neighboring DNA ends simultaneously, which somewhat decreased the successful ligation rate. To date, ChIA-PET has not been used at the single-cell resolution by any researchers.

Lin developed a new method, called DLO HiC, to solve this problem^4,5^. The method involves inserting two linkers with a specific sequence between neighboring DNA fragments. These linkers contain a specific sequence with digestion recognition sites at the junction position. After successful ligation, MDA amplification can be performed, and the ligated junctions can be enriched by specific digestion with recognition sites on the linkers. This technique relies on the adapter’s specific sequence instead of biotins for enrichment, which reduces the loss of enrichment in minimal input DNA. Multiple displacement amplification (MDA) can generate ample DNAs for downstream enrichment, eliminating the need for biotin purification and amplifying the DNA before enrichment. However, the successful ligation rate may decrease because this step requires two linker ligations in the junction position, meaning that the fragments must be ligated three times.

While enrichment-based methods have their place, alternative approaches like Dip-C^6^, and multiomics scCARE-seq^7^, HIRES^8^, sn-m3C-seq^9^ directly link neighboring fragments, bypassing the need for end repair. This offers the advantage of a more efficient sticky-end ligation in contrast to the methods employing biotin-incorporated blunt ends. Nevertheless, it’s important to be aware that full genome sequencing might lead to around 90% of reads being discarded, which could influence the detection of rare DNA conformations.

When carrying out single-cell chromatin conformation capture (3C) experiments, the selection of methods merits careful consideration. In order to cut down on sequencing costs and detect rare contacts more effectively, a method that can manage large-scale cell numbers and ultra deep sequencing per cell is required. Our approach, HiChew (High efficient chromatin conformation capture without pre-enrichment), is notable for its high efficiency. HiChew is unique as it circumvents the need for pre-enrichment prior to amplification. We employ a high-efficiency sticky end ligation method, akin to Dip-C and HIRES, which reduces the steps required before amplification. Upon amplification, we mark the ligation scar motifs with corresponding methyltransferase and enrich the ligation scars using a methylation antibody. In summary, our approach has delivered the greatest efficiency for single-cell 3C to date, enhancing not only single-cell 3C but also simplifying conventional Hi-C in common labs.

## Results

### Principle of the HiChew design

Our HiChew design process simplifies down to four key steps before PCR. First, we crosslink the DNA strands and proteins together by formaldehyde. Next, we treat the DNA on the GATC motif using DpnII. Then, we connect nearby digested DNA ends, leaving behind a GATC ligation scar – this is based on the principle of proximal ligation. After cell breakdown, we can choose either a adaptor ligation-based or Tn5-based DNA library construction method, depending on the amount of DNA and your specific requirements (Fig. 1A).

**Fig.1.**
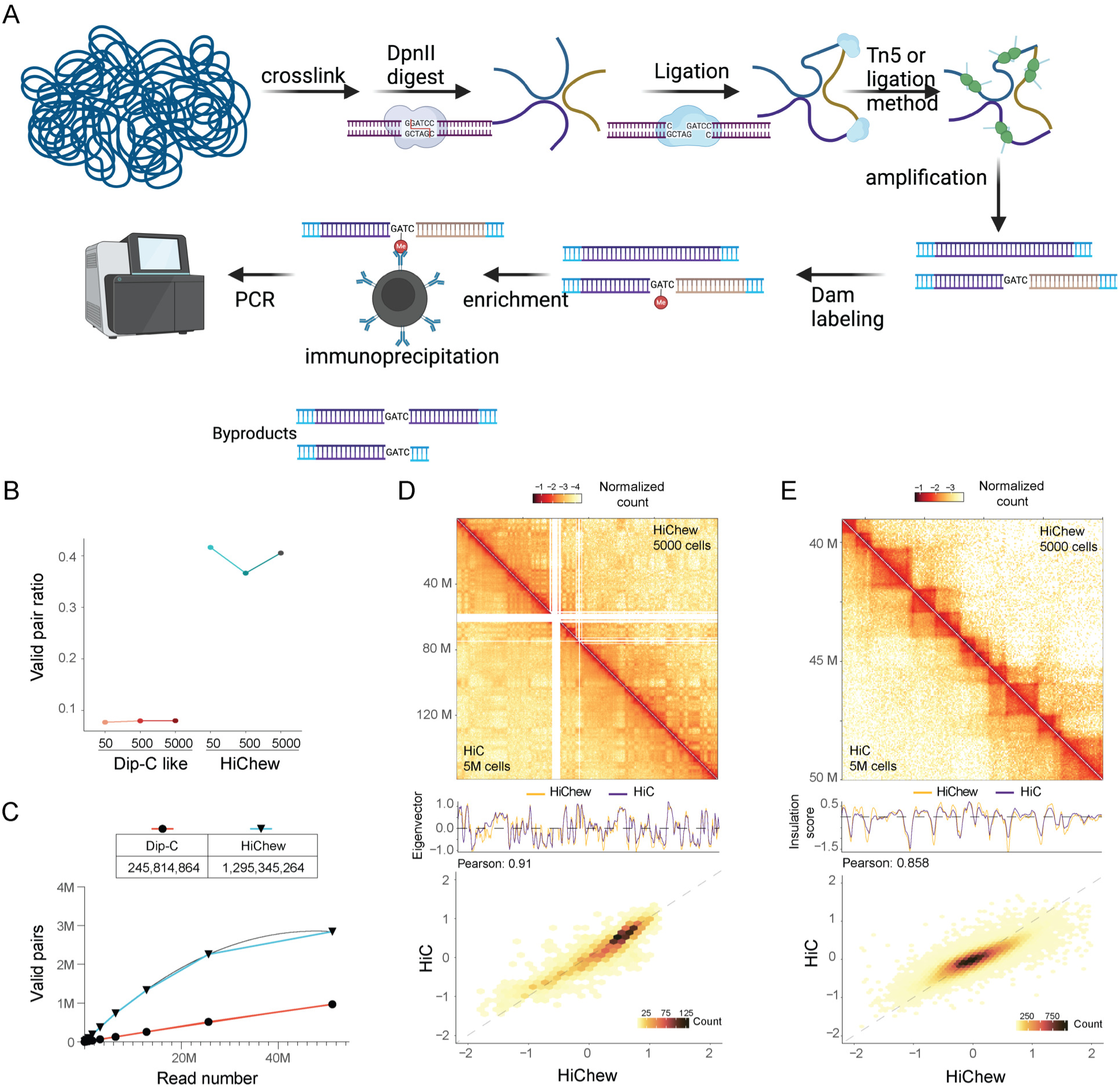
Overview of the HiChew experiment procedure and evaluation. (A) A simple diagram of the experimental procedure. (B) The valid pair ratio of HiChew and Dip-C with different cell inputs. (C) The saturation curve of the Dip-C and HiChew for 50 cells. (D) The top panel displays the eigenvalue plot for HiChew and Dip-C on chr7. The middle section compares chromatin contact maps of Hi-C with HiChew. The last panel shows the eigenvalue correlation between Hi-C and HiChew. (E) The top panel shows the insulation score plot of HiChew and Dip-C on region Chr7:39M-50M. The middle section again compares chromatin contact maps of Hi-C with HiChew. The bottom panel shows the insulation score correlation between Hi-C and HiChew.d

Once we’ve effectively copied the DNA through PCR amplification, we use the dam methyltransferase to mark the ligation scar with m6A methylation on the GATC motif. After this, we apply the immunoprecipitation (IP) protocol to enrich the ligation scars using m6A antibody pull down (Fig. 1A).

This approach efficiently solves the problem of low cell quantity or single-cell Hi-C biotin enrichment. The process omits steps such as end repairing, biotin incorporation, or adapter insertion, focusing instead on digestion and direct ligation. It boasts high ligation efficiency, minimizing steps before PCR amplification to lessen potential cell or DNA loss. The superior post PCR enrichment process enables us to conduct the enrichment with ample DNA, mitigating the loss of biotin enrichment before PCR.

This leads to several questions. Does GATC only represent ligation scar patterns? There are two types of GATC motifs that may not necessarily indicate ligation scars.

Firstly, there are GATCs that cannot be broken down by DpnII. These undigested GATCs could be due to less effective digestion within the cell, possibly triggered by crosslinking or specific chromatin structures. We compared in-cell digestion with purified genome digestion and found that only about 30% of GATCs remained undigested. This suggests that digestion efficiency could significantly affect our capture scar efficiency, a hypothesis confirmed by our subsequent data analysis.

Secondly, there are GATC sequences that have been digested but not ligated. These sequences often appear at DNA strand ends, making them easy to eliminate. Therefore, we are not overly concerned about these two byproducts.

Generally, the HiChew method aims to improve the efficacy of chromatin conformation capture experiments. This is particularly beneficial in single-cell experiments, where enrichment can lead to DNA loss. The process involves capturing ligation scars after amplification using the GATC motif, which the enzyme dam methyltransferase recognizes.

### The performance of HiChew is similar to Hi-C in low input bulk sample

We initially tested biotin-based Hi-C, unenriched Dip-C like method, and HiChew using different cell quantities: 5000, 500 and 50 cells. The libraries were sequenced using 100∼200 million paired-end reads. To compare the results, we used the gold standard Hi-C from Rao (5million cells). Our initial analysis focused on evaluating the effectiveness of these methods.

In our labs, we tried the standard Hi-C protocol on 5000-50 cells but, unfortunately, we didn’t obtain any significant signals. This might be because the low input cells didn’t meet the standard HiC requirements. We then turned to traditional high-input Hi-C data that we had downloaded, which used biotin enrichment, and found it had a valid pair ratio of 95% (452M valid pairs out of 473M reported pairs in 5 million cells) (valid pair ratio is equal to valid pair/aligned reads, details in methods). On the flip side, HiChew gave us a valid pair ratio of around 42% (44M valid pairs out of 106M reported pairs in 5000 cells) (Fig. 1B), which is not too far off from Hi-C. The unenriched method, otherwise known as “Dip-C like”, only managed to reach a validation rate of 7% (Fig. 1B). This means HiChew is approximately 6 times more efficient at capturing valid pairs than Dip-C. Something worth noting is that HiChew had roughly 26% undigested GATC sites, a byproduct that wasn’t efficiently processed during in-cell digestion according to our sequencing data. This might explain why the number of valid pairs is lower compared to Hi-C. Additionally, the lower enrichment ratio could be due to the antibody’s reduced affinity compared to the biotin-streptavidin binding. In conclusion, HiChew, by using post-PCR enrichment, manages to reach an efficiency level that’s comparable to traditional Hi-C.

We are exploring the possibility that an increase in capture efficiency could potentially compromise the sensitivity of these technologies, resulting in lost of contact information. A limitation of single-cell/low input Hi-C is that while biotin purification can enrich valid pairs and decrease sequencing costs, it also leads to a significant loss of DNA and decreased signal, making it challenging to observe patterns even with an input of 5000 cells (Sfig. 1A). This is likely why most recent technologies are leaning towards Dip-C for single cell studies.

Our sequencing saturation test shows that HiChew, using 50 cells, gives a higher estimated maximum contact (1295M) compared to Dip-C with the same number of cells (245M) (Fig. 1C). This implies that HiChew can extract five times more contact information from a limited number of cells than Dip-C, indicating a higher sensitivity. This makes sense as the processes before PCR are the same for both methods. Once we move past PCR amplification, when there’s ample DNA, any loss in the enrichment steps becomes less of a concern. The post-PCR enrichment of HiChew notably boosts the valid contact yield, reduces junk sequences, and improves sensitivity in contact detection. In each sequencing depth, HiChew could provide 4-6x more valid contact than unenriched Dip-C (Fig. 1C). To sum it up, HiChew matches the enrichment efficiency of Hi-C, even with 10-100 times fewer cells. It also exceeds Dip-C’s capture capability while generating data more efficiently and requiring fewer steps in the process.

We’ve performed additional quality tests on HiChew to ensure the accuracy of its data generation in chromatin conformation capture. We also looked into how different input cells affect the final results. In terms of the cis-trans ratio, HiChew and similar methods like Dip-C all produced comparable outcomes (Sfig. 1B). From 50 to 5000 cells, the distance decay curve remained consistent across all methods (Sfig. 1C). Notably, HiChew, when used with 5000 cells, produced results on par with Hi-C in multiple resolutions, including AB compartment(Fig. 1D), TAD insulation (Fig. 1E), and loops (Sfig. 1D). However, we observed a decrease in similarity among AB compartment scores and TAD insulation scores as the number of cells diminished (Sfig. 2, Sfig. 1D). It’s important to highlight that the intricacy of structures was influenced by the number of input cells, considering each cell possesses a unique chromatin conformation structure. Therefore, a larger cell count is likely to provide a more representative view of the bulk cell phenomena. When comparing performance across all scales and cell inputs, Dip-C fell short of HiChew due to its lower yield of valid contacts (Fig. 1DE, Sfig. 2BCDEF). An alternative approach involves digesting the genome with AluI and labeling the ligation scar using AluI methyltransferase. This method also showed a high correlation with DpnII batches, demonstrating the robustness of the technology (Sfig. 2A). Overall, HiChew successfully generated chromatin conformation with a notable enrichment for smaller input cell numbers.

### The high throughput single nuclei HiChew (snHiChew)

HiChew stands out as a sensitive technique for accurately detecting chromatin conformation, even with a small number of cells. Understanding the changes in chromatin conformation in individual cells is key to our broader scientific knowledge. Other methods, like Dip-C, HiRES, and scCARE-seq, also use the unenrichment approach to sequence the whole genome. This helps to avoid losing contacts during biotin incorporation and purification stages. As a result, we’ve adjusted HiChew to effectively capture chromatin conformation in single cells without pre-enrichment. This has led to snHiChew, a high-throughput and efficient method focused on single nuclei.

The snHiChew method takes inspiration from HiChew, involving similar steps. We start with crosslinking, then digest the DNA and ligate it into concatemers to capture chromatin conformation. Next, we employ HpyCH4V to digest the DNA concatemer, preparing it for barcode attachment. This is followed by dA-tailing and ligation with the first round of barcoded adaptors (Fig. 2A).

**Fig.2.**
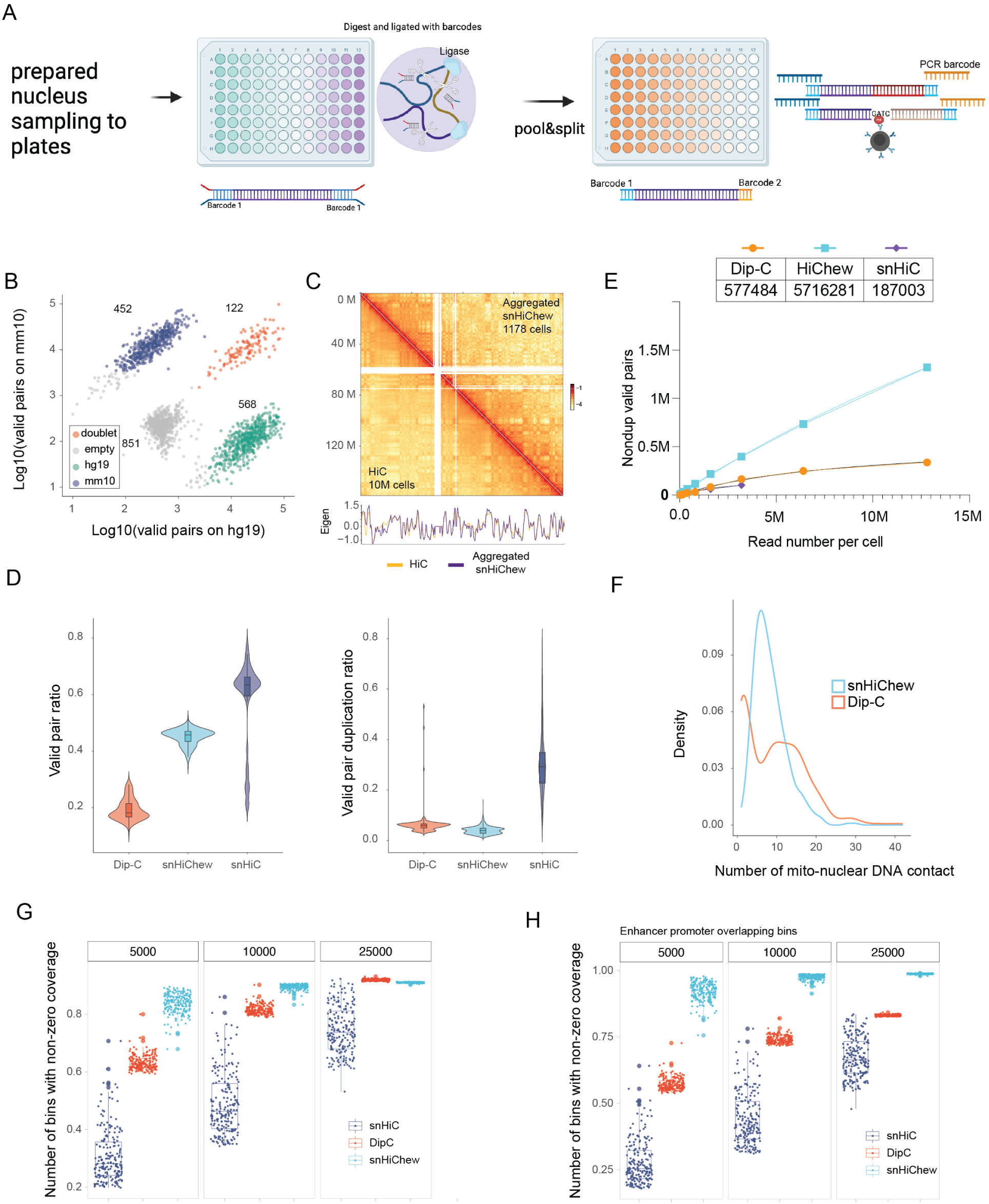
The snHiChew experiment procedure and evaluation. (A) The snHiChew process is illustrated. (B) We used a two-cell mixture, 3T3 (mouse) and HEK293T (human). The valid contacts in each cell are summarized, aligned to either the mouse (mm10, y-axis) or human (hg19, x-axis) genomes. (C) The top panel shows the eigenvalue from Hi-C (326M valid pairs) and from HiChew (combined 1178 cells, 326M valid pairs). The bottom panel reveals the contact maps on chr7. (D) The valid pair ratio is calculated by the count of valid reported pairs over the count of reported reads. The pair duplication ratio is calculated by duplicated pairs over valid pairs. Valid pair per read is calculated by the count of deduped valid pairs over total reads. (E) The valid pair saturation curve is displayed. The x-axis represents the sequencing depth threshold for each cell, and the y-axis represents the log10(deduped valid pairs) from the corresponding sequencing depth. (F) The density of false positive ligation counts between chromosome and mitochondrial DNA is depicted. (G) The genome was divided into sizes of 5000bp, 10000bp, 25000bp. The count of bins with nonzero contacts in each cell was summarized and divided by the total bins. (H) The genome was divided into sizes of 5000bp, 10000bp, 25000bp, with selection only for bins that include an enhancer or promoter. The count of bins with nonzero contact in each cell was summarized.

At this phase, we gather the nuclei and segregate them into various groups on the secondary plates. We then conduct PCR to affix the secondary barcode (Fig. 2A). Utilizing the pool-split method, we theoretically have the capacity to produce 147,456 barcodes (384×384), potentially accommodating up to 20,000 cells at a 0.13 duplex rate. In a smaller batch test, a 98×98 barcode system could potentially handle around 2000 cells.

After PCR amplification of the entire genome library, we utilize the methyltransferase to mark the junction scars with m6A. Then, we employ the m6A antibody to enrich the valid pair reads (Fig. 2A). Worth noting is that we should steer clear of the GATC motif when designing the barcode. This is to avoid being targeted by the methyltransferase, which could potentially skew the data.

In our efforts to guarantee snHiChew data’s precision and dependability for single cell identification, we initiated our experiments with a combination of HEK293T and 3T3 cells. The successful detection of human and mouse reads in snHiChew underscores its efficiency in identifying individual cells - we identified 568 human cells and 452 mouse cells from an expected 1200 (Fig. 2B). We also observed 122 doublets, aligning with a 0.1 collision rate (Fig. 2B). Utilizing UMAP clustering grounded on the chromatin conformation structure, we could easily distinguish between these two cell types (Sfig. 3A).

We also conducted deep sequencing on another batch of HEK cells that fulfilled strict quality standards, resulting in an average of 500,000 chromatin contacts per cell (Sfig. 3B). To gauge snHiChew’s accuracy, we carried out an aggregated pseudo-bulk analysis (1178 cells) for quality control. As expected, we found that the grouping of similar cell reads accurately reflected the bulk Hi-C chromatin structure in terms of distance decay curve, AB compartments, TAD insulation and loop analysis (Fig. 2C, Sfig. 3C-H,F-G).

Next, we conducted a comparison between snHiChew and the well-known Dip-C, representing the unenrichment method, together with snHi-C, which symbolizes the biotin enrichment method. We then evaluated the data usage, false positive rate, and resolution, which are crucial factors in Hi-C experiments.

In our quest for data usage efficiency, we first normalized all cells to 100,000 reads per cell. Dip-C and snHiC-seq yielded valid pair ratios of approximately 15% and 65%, respectively, while HiChew offered around 48% (Fig. 2D). As anticipated, snHi-C, which uses biotin purification, showed the highest data usage efficiency, despite significant DNA loss. To support this, it’s worth noting that the duplication rate can generally provide insights into library capacity and retained DNA information. Here, snHi-C had a notably higher duplication rate, about four to five times that of HiChew and Dip-C (Fig. 2D). This suggests a lower library capacity for snHi-C, leading to fewer contacts captured and quicker saturation (Fig. 2E). Lastly, to examine cost efficiency, we looked at the valid pair per sequence read. After removing duplicate reads, HiChew consistently had the highest valid pair in each sequencing depth, still three to seven times that of Dip-C (Fig. 2E). This suggests that HiChew can produce three times more unique valid contacts within the same budget.

In our assessment of false positives, we employed the cis-trans ratio, a consistent measure across all methods (Sfig. 3J). To boost our ability to spot false positives, we included a quantification step of interactions between nuclear DNA and mitochondria DNA. HiChew demonstrated a significant decrease in false positives when compared to Dip-C (Fig. 2F). This improvement is thanks to the enrichment of the ligation scar, which in turn boosts specificity.

We’ve further analyzed the maximum resolution of a single cell, which is typically determined by the maximum contacts per cell. Our contact saturation analysis showed snHi-C-seq reaching saturation quickly with only 187K maximum contacts (Fig. 2E). The percentage of valid pairs then dropped precipitously (Sfig. 3I), suggesting that snHi-C-seq may not have the sufficient sensitivity to capture sufficient valid chromatin interactions. This underperformance can largely be attributed to DNA loss during purification.

Contrastingly, HiChew outshined the rest by delivering an estimated maximum of 5716K contacts per cell, over ten times more than Dip-C. This suggests that HiChew could provide the most comprehensive structural information. We were able to generate between 500K and 2M contacts per cell, marking the highest resolution in single cell chromatin contact maps to date. We’re optimistic about the potential of HiChew technology to deliver even higher contacts per cell.

When comparing various methods, snHiChew proved to have a superior resolution at approximately a 5kb bin with 90% coverage (Fig. 2G). Our high resolution has allowed us to identify the greatest number of enhancer-promoter interactions from single cells compared to other technologies (Fig. 2H). Notably, we achieved this resolution with a substantially smaller sequencing budget, without resorting to a blunt increase in sequencing depth. This could significantly enhance our understanding of transcription regulation synergy within cells.

In terms of data usage efficiency, maximum contacts captured, and fine resolution, snHiChew has proven to be more efficient than existing methods. Our comprehensive validations and benchmarks affirm the high-quality data produced by snHiChew, with high resolution contact map per cell also observed.

### subTADs melting with specific gene activation in B compartments

Chromatin segregates into two large compartments: A and B^10^. Generally, heterochromatins reside in the B compartment, close to the lamin area, which is related to silent expression. However, previous research has discovered that genes in the B compartment can still express in the cell. The method of their expression remains a mystery in the 3D genome area.

Our study identified numerous long coding genes situated within the B compartment (Sfig. 4A). We chose a typical region, spanning from 145Mb to 155Mb, as a representative example (Fig. 3A). The stretch from 145MB to 148MB, encompassing the long gene CNTNAP2 with a genomic size of 1.5MB, was located within the B compartment and covered by 3 subTADs (blue dash). The subsequent region, extending from 148MB to 152.5MB, represented another A compartment containing over 30 genes. The final section, from 152.5 MB to 155MB, was again within the B compartment. The insulation score showed three drops in the segment from 145MB-148MB, with the presence of CTCF peaks on the subTAD boundary further confirming the existence of the subTAD (orange arrow). Interestingly, we observed some active histone modifications, such as H3K4me3 signal on the CNTNAP2 promoter and H3K27ac on the gene body. Furthermore, we observed the weak RNA expression of CNTNAP2. This presents a paradoxical scenario, considering that subTAD segregates the self-domain and insulates the interactions between TADs, raising the question of how transcription could pass through these domains.

**Fig.3.**
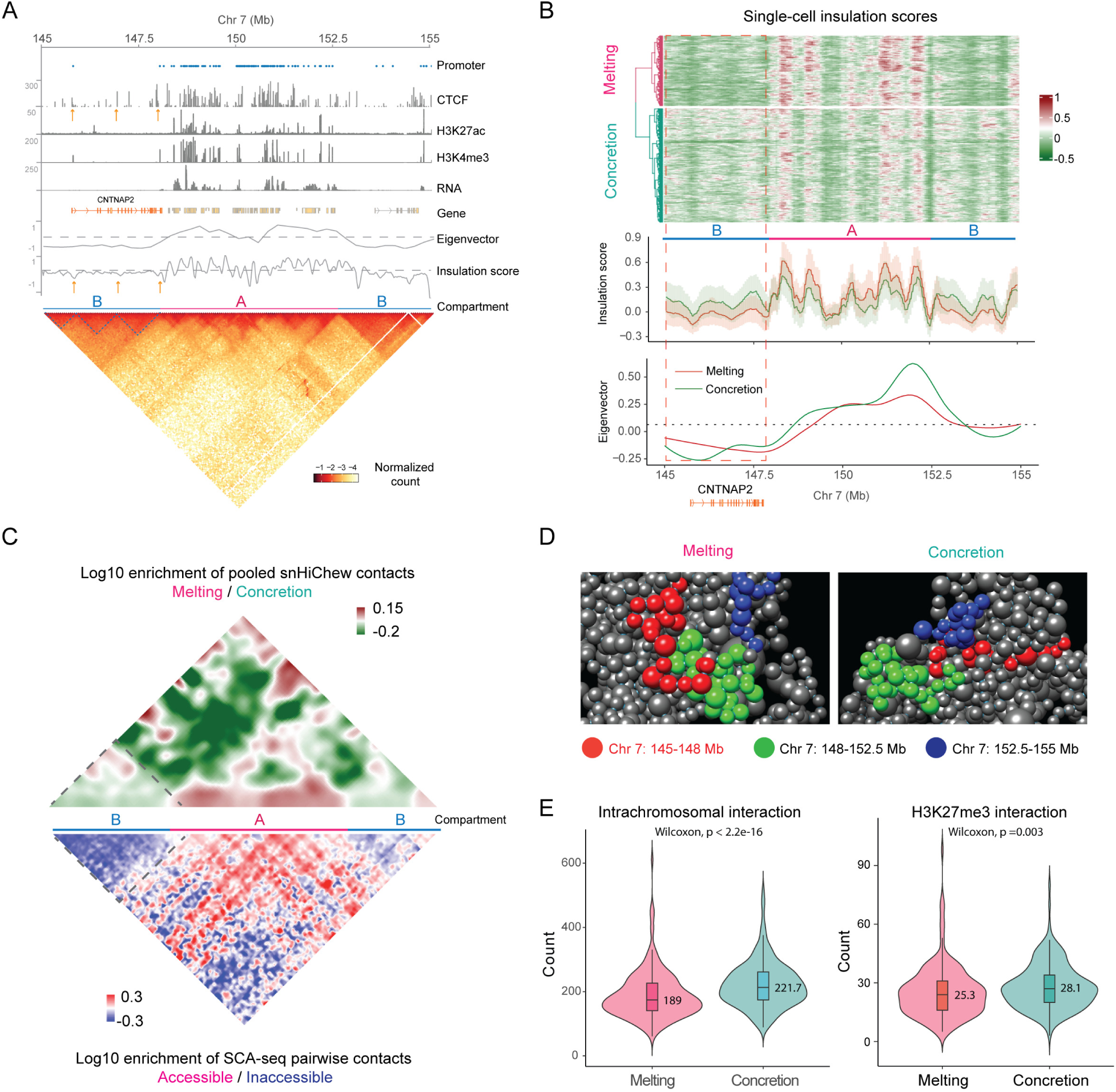
Changes in Gene Expression Relating to Insulation. (A) Here’s an example from the 145-157MB region. We used ChIP-seq data to get CTCF, H3K27ac, and H3K4me3 peaks. We calculated Eigen value and Insulation score from the contact maps. ’Promoters’ points out where each gene’s promoter is located. We used software to identify Compartment AB. The bottom heatmap provides a view of the chromatin contact maps. (B) This panel shows how we clustered the insulation score for individual cells. The bottom part demonstrates the mean insulation score and eigen value of the melting and concretion cluster. (C) The top part displays the contraction heatmaps of the melting cluster contact density compared to the concretion cluster contact density (melting/concretion). The bottom part displays the contraction heatmaps of the accessible chromatin contact density contrasted with the inaccessible chromatin contact density (accessible/inaccessible). (D) This panel gives you a 3D model of the melting status and concretion status. The red balls stand for the 145-148M region containing the CNTNAP2 genes, the green ball for 148M-152.5M and the blue balls for the 152.5M-155M. (E) The violin plot illustrates the intra-chromosome and H3K27me3 interaction frequency with the 145-148M regions in individual cells.

Our initial hypothesis is that if transcription occurs throughout this entire gene, it could potentially loose the DNA and dissolve the segregated subTADs. To investigate this, we first attempted to identify cells with activated and silenced CNTNAP2. We performed a single-cell cluster analysis on HEK293T cells, taking into account the eigenvalue (representing the AB compartment) and the insulation score (representing TADs). Despite the high homogeneity of HEK293T cells, the abundance of chromatin contacts allowed us to easily sort the cells into two distinct clusters based on the insulation score (Fig. 3B). One cluster showed a decrease in insulation score in the B compartment (red dash box) and an increase in the adjacent A compartment, indicating a dissolution status (melting label) in the 145Mb-148Mb B compartment. Conversely, the other cluster showed opposite trends, suggesting a solidification status (concretion label) in the 145-148MB B compartment. Comparing the two clusters, we noted a variance in the B compartment insulation score from −0.01 to 0.11 (Fig. 3B insulation, Sfig. 4C), indicating possible subTAD dissolution. However, this classification becomes less clear when valid contacts per cell fall below 250k (Sfig. 5).

Looking at the two status cells (Fig. 3C), it’s evident that the concretion status cells form more contacts within the three subTADs (green area under the dashed triangle), as shown by the contact comparison. In contrast, the melting status cells form more contacts outside these subTADs (red area under triangle). This could indicate that the subTADs might merge into a larger TAD. The decrease in the insulation score in this area supports this idea (Fig. 3B, Sfig. 4C). It’s plausible that this process could be a way to remove obstacles in the RNA polymerase II path, potentially related to CNTNAP2 transcription. To reinforce this gene activation, we calculated the eigenvalue for each cluster. While the eigenvalue didn’t reveal a significant difference between the two clusters, the melting status did show a slight increase in eigenvalue compared to the concretion status in the 145-148MB B compartment, suggesting that the gene may move slightly for expression (Fig. 3B eigen, Sfig. 4B). This idea is also backed by recent findings that B compartment gene expression doesn’t significantly impact the B compartment eigenvalue^11^.

To further explore the relationship between gene expression and chromatin conformation, we’re leveraging SCA-seq^12^, our multiomics chromatin conformation capture technology. This allows us to examine changes in chromatin accessibility and genome structure at a single molecule resolution. We’re comparing the interactions of accessible and inaccessible chromatin domains. Interestingly, we’ve observed that inaccessible chromatin(blue) tends to form smaller subTADs near the diagonal line, while accessible chromatin (red) extends into larger TAD areas in the B compartment (Fig. 3C, bottom panel). This distribution mirrors the patterns of melting and concretion clusters, hinting at a possible association between the melting cluster and accessible chromatin. In the A compartment, accessible interactions align closer to the diagonal line, with a corresponding enrichment of melting status in the same area. Consistent with previous research, chromatin accessibility is strongly linked with active histone markers and transcription activity. This strengthens the hypothesis that the dissolution of subTADs in the B compartment might be intrinsically linked to chromatin accessibility and gene activation.

Apart from the regions mentioned, we’ve identified several others with exceptionally long genes (>1MB). These regions also exhibit high insulation within the gene body (insulated score>0.1) and show a similar pattern of subTAD dissolution (Sfig. 6). Across the board, our findings indicate a substantial link between subTAD dissolution within the gene body and gene activation.

### 3D modeling explain the melting movement

In our research, we employed 3D modeling to examine the changes in chromatin folding between two cellular states (Fig. 3D). We observed that in the concretion state, the B compartment (red balls) was densely packed and in close proximity with the rest of the genome. On the other hand, in the melting state, the 145-148MB B compartment was loosely organized and situated on the chromosome surface. This less structured arrangement aligns with our finding of a lower insulation score, adding further support to our results. We then proceeded to explore whether these movements were associated with the regulation of specific epigenetic markers.

Our research primarily aimed to find out if changes in chromatin folding could trigger interactions with active histone markers H3K4me3 and H3k27ac. Yet, we didn’t see any noteworthy shifts. Instead, we found that transitioning from a concretion to a melting state considerably decreased intrachromasomal interactions (Fig. 3E). This supports our 3D model suggesting a packing with other chromatin in the B compartment’s concretion state. Upon further investigation, we found that interactions between the silenced H3K27me3 marked region and CNTNAP2 significantly dropped in frequency during the melting state (Fig. 3E). These insights imply that the melting of the subTAD in the B compartment is tied to the gene’s relocation to the surface of the chromosome, moving it away from the H3K27me3 dominated region. This observation aligns with the Seq-Fish finding that most gene expression occurs on the chromosome surface^13^ and with the shifting of the B compartment gene in the expression^11^.

### Insulation Dynamics in Relation to Gene Bursting

Once we comprehended the mechanism of long gene activation in the B compartment, we organized the cells according to the replication score. This score provides detailed insight into the cell replication cycle from its early to late stages, and aids in observing the gene activation sequence rules. Interestingly, we observed dynamic changes in the insulation score as the cell progressed, corroborating previous research findings (Fig. 4E)^14,15^.

**Fig.4.**
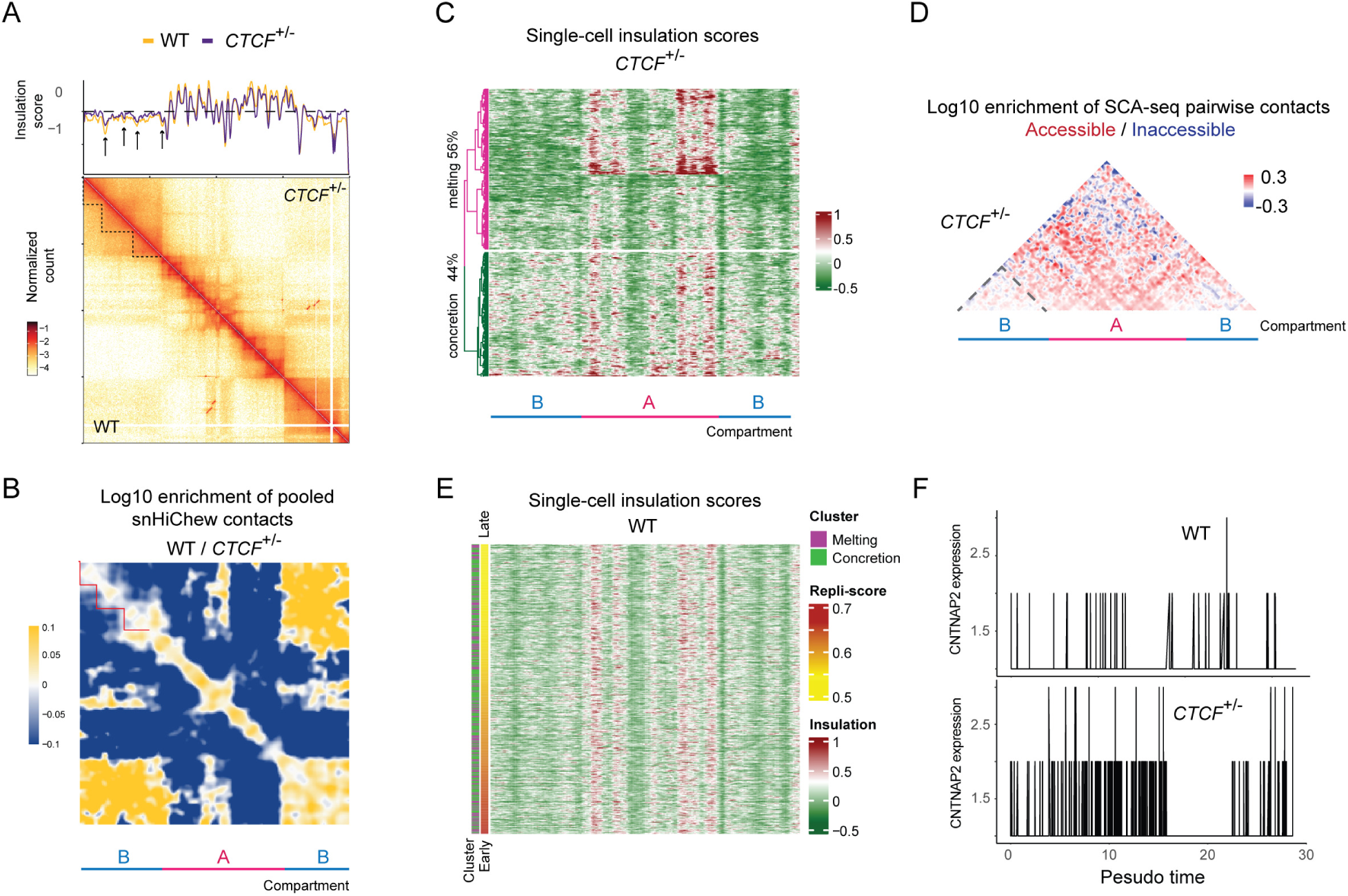
CTCF+/- results in decreased insulation. (A) Chromatin contact map from 145MB to 155MB of the WT and CTCF+/-. The upper panel displays the insulation score of this region in WT and CTCF+/- cells. (B) Comparison heatmap of contact density between WT and CTCF+/-. (C) Cluster analysis of insulation scores in single cells of CTCF+/-. (D) Comparison heatmap of contact density between accessible and inaccessible chromatin in SCA-seq. (E) Insulation scores of single cells, sorted by replication score. Cells from melting and concretion clusters are marked with purple and green bars, respectively. (F) Expression of CNTNAP2 in single cells, cells are arranged by pseudo time calculated via RNA velocity.

In the heatmaps, we identified the melting clusters within the single cell insulation scores organized with the replication score. These clusters sporadically appear as the cell cycle continues, potentially suggesting an intermittent expression of the CNTNAP2 gene. To confirm these observations, we further examined gene expression on a single cell level. Our study found that CNTNAP2 displayed corresponding intermittent expressions (Fig. 4F, WT), following the same pattern. This supports past research indicating that the expression of this gene behaves like a switch, toggling between activation and silencing while coinciding with dynamic changes in TAD.

### CTCF maintain the synergic interaction in cells

Our latest observations show that CTCF is typically located between TADs and even subTADs in the b compartment, as supported by previous research ^16,17^ (Fig. 3A). This interesting discovery led us to investigate if a decrease in CTCF protein expression could perhaps increase the melting frequency.

In our study to understand the effects of reduced CTCF on entire cell cycles, we used CTCF +/- cell lines with significantly lower CTCF protein expression (Sfig. 8). By sequencing 1020 CTCF+/- cells through snHiChew, we managed to achieve an average of 500k valid pairs per cell. Our contact maps of cell aggregation revealed a slightly weaker subTAD in the 145-148MB B compartment compared to the wild type (Fig. 4A). When normalized for comparison, the CTCF+/- cells (blue) showed a more noticeable enrichment in the large TAD area, while the wild type (yellow) had more enrichment in the subTADs. This suggests a looser compaction in CTCF+/- (Fig. 4B).

Our analysis of insulation revealed a less pronounced drop signal at the subTAD boundary in CTCF+/- (see Fig. 4A for insulation). This suggests that these subTADs have less defined boundaries. In terms of single-cell changes, they were notable. For instance, the number of cells in the melting cluster jumped from 30% to 56% (Fig. 3B, Fig. 4C). This suggests that the melting frequency during the cycle has increased. Given the connection between the melting process and chromatin accessibility in the B compartment, we used SCA-seq on CTCF+/- cells to examine any changes in chromatin accessibility on the contact map. When comparing SCA-seq between CTCF+/- (Fig. 4D) and the WT(Fig. 3C), we observed a significant increase of open chromatin on the CNTNAP2 area (Fig. 4D). This suggests that a higher melting frequency might be linked to enhanced chromatin accessibility, possibly affecting gene expression.

Our analysis of single-cell RNA seq data^18^ revealed a significant rise in average CNTNP2 expression due to the lack of the subTAD insulating the genes (Fig. 4F). When we arranged the cells as pseudo times, we noticed an uptick in sporadic frequency of CNTNAP2 expression in CTCF+/- cells (Fig. 4F). This observation corresponds with an increase in melting frequency. In other regions (Sfig. 7A), we also found the melting status increased frequency and the globally long gene expression increased (Sfig. 7B). It highlights the essential role of CTCF and insulation in suppressing long gene expression in the B compartment. These insights indicate that CTCF plays a part in maintaining the subTAD, which aids in insulating long gene expression.

Overall, the study introduces HiChew, a technique that effectively captures chromatin conformation in low-input cells and is even suitable for single-cell applications. HiChew proves to be as efficient as traditional Hi-C techniques, laying the foundation for snHiChew, which outperforms in efficiency and resolution. The team noted changes in chromatin conformation during gene activation, finding a substantial link between the dissolution of subTAD and gene activation. They further explore the role of CTCF in preserving subTAD and its impact on gene expression isolation.

## Discussion

In this study, we introduced highly efficient technologies, HiChew and snHiChew, which significantly address a major issue in this field: the inefficient incorporation of biotin enrichment into single cell analysis, resulting in significant loss of valid contacts. Traditionally, many researchers have resorted to Dip-C that rather crudely sequences the whole genome without enrichment, leading to a waste of approximately 92% of the data. This approach, while enabling high sensitivity to capture more interactions, has dominated the field for many years.

Previously, the idea of enriching the ligated fragments after PCR, which could generate valid contacts without sacrificing DNA loss, was not widely explored. Our method, HiChew, solves this long-standing problem and is of increasing importance in current studies.

The latest studies ^19,20^ underscore the dynamic nature of chromatin conformation, often requiring a larger sample size of up to 100,000 cells. Given that each cell needs 2M reads to generate 200K valid contacts, this translates to a need for sequencing 200,000M data, which comes with a significant budgetary requirement given the current cost of approximately $7/MB. In our work, we have demonstrated that over 500K contacts per cell can uncover more phenomena. Additionally, we found that methods not utilizing enrichment have a higher false positive rate, which could impact the quality of data from shallow sequencing in single cell Hi-C experiments. As research continues and the need for more cells grows, the expense and waste associated with sequencing will increasingly become a critical concern.

Our methodology does have some limitations. Specifically, we’re confined to choosing the restriction enzyme that pairs with the methyltransferase for marking the ligation scar, such as DpnII-dam methyltransferase and AluI-AluI methyltransferase. In our research, we employed DpnII and AluI to evaluate the abilities of HiChew. It’s important to recognize that the commercial availability of these enzymes is on the rise, with roughly 20 different enzymes currently on the market. This could broaden the application of our methodology to a range of conditions. Another aspect to ponder is whether the resolution of HiChew can be improved by utilizing multiple enzyme digestion. The theoretical resolution of the DpnII digestion is approximately ∼700bp. If our goal is to break down genome into smaller fragments for better resolution, we might think about using a blend of multiple enzymes, such as a combination of DpnII and AluI. This could theoretically enhance the resolution to ∼400bp, albeit at the cost of some sensitivity.

In our research, we employed deep sequencing and high-resolution snHiChew to find that as the insulation score of compartment A increased, that of compartment B 145-148MB decreased. This raises the question: is the gene expression in compartment A halted with an increase in insulation? This is a complex issue. However, our data suggests that the increase in insulation in compartment A is accompanied by an increase in chromatin accessibility. Comparisons between accessible and inaccessible chromatin support this hypothesis, with accessible chromatin segregating around the diagonal line. The subsequent question to consider is whether segregation in compartment A assists neighboring compartments in converging. This is a possibility, given that similar hypotheses have been proposed through SPRITE technologies^11^. Our SCA-seq data also indicates that accessible chromatin is enriched in the subcompartment region between two neighboring B compartments. However, no significant proximity of B compartments was observed during the insulation melting process in our single-cell 3D modeling. In contrast, we found that melting B compartments could potentially be relocated from inside to outside. This correlates with previous publications that demonstrated the most active genes on the chromosome surface through Seq-FISH. The minor position change of this compartment could align with these previous findings.

In our experiments involving CTCF knockdown, we noticed that this process affects subTAD insulation on an individual cell level and is linked with the frequency of gene expression. Similar observations have been made by other specialists in this field, who discovered that altering the CTCF motif increases the frequency of interaction between promoters and other enhancers^21^. As a result, a decrease in insulation within the subTAD could either be a consequence or cause of extensive gene expression in the B compartment. Although we can’t dismiss the possibility that the increase in interaction with nearby enhancers encourages gene expression, or that the removal of a silencer triggers gene expression, these possibilities extend beyond the scope of this methodology study.

In our labs, we’ve effectively utilized snHiChew in animal tissues, such as testis, where it maintained stable performance. Furthermore, HiChew technology can be incorporated into ChIA-PET^17^ and HiChIP^22^, which typically require a significant number of cells. With HiChew’s assistance, HiChIP only needs a single immunoprecipitation, simplifying the process and increasing the success rate while also reducing the number of cells required. This could significantly streamline these complex technologies. Additionally, this technology can be integrated with other multi-omic technologies like HIRES (RNA+HiC) and Methyl-HiC (methylation+HiC). Overall, HiChew represents an efficient approach to chromatin conformation studies.

## Method

### Cell Type and Cell Culture

Mouse embryonic fibroblasts cell line 3T3 was obtained from ATCC. Human embryonic kidney cell line Hek293T and CTCF+/- Knockout 293T Cell Line were obtained from Abclonal Technology Co.,Ltd. 3T3, Hek293T and CTCF+/- Knockout 293T were cultured in DMEM (Gibco, no. 11965092) medium supplemented with10% FBS (Gibco, no. 10099141) and 1% penicillin–streptomycin (Gibco no. 15140122), under the conditions of 37°C, 5% CO2.

### Cell Sample Fixation

First, remove the medium from the culture dish. Then, add 3 ml of room temperature 1×PBS to gently rinse the cells, and aspirate the PBS. Add 2 ml of trypsin to the 10 cm culture dish. Gently shake the dish to evenly distribute the trypsin across the entire cell layer. Place the culture dish in the 37 °C incubator for digestion.

Next, add 4 ml of DMEM complete medium and pipette several times to detach the cells from the dish. Transfer the cells to a 15 ml centrifuge tube and centrifuge at 400g for 3 minutes. Discard the supernatant and wash the cells twice with 1×PBS. After centrifuging at 400g for 3 minutes, resuspend the cells with 2 ml of 1×PBS.

Add 500 μl of 2% formaldehyde (16% wt/vol formaldehyde ampules, Life Technologies, no. 28908, prepared in 1×PBS) to 500 μl of resuspended cells for a final concentration of 1% and fix at room temperature for 10 minutes. Then, add 50 μl of 2.5 M glycine, terminate the fixation at room temperature for 5 minutes, and centrifuge at 500g for 5 minutes. Remove the supernatant and wash twice with 1×PBS.

Finally, resuspend the cells in 1 ml of a 1×PBS solution containing 3mM EGS (Thermo, no. 21565) and fix at room temperature for 45 minutes. Add 200 μl of 2.5 M glycine and terminate the fixation at room temperature for 5 minutes. Centrifuge at 500g for 5 minutes, remove the supernatant and process to cell lysis or flash-frozen in liquid nitrogen.

### Cell Lysis

Resuspend the cell pellet in 500 μl of membrane permeabilization buffer (10 mM Tris-HCl (pH8.0), 10mM NaCl, 0.2% CA630, 1× proteinase inhibitors cocktail). Then, incubate it on ice for 10 minutes. After that, centrifuge at 500g at 4°C for 3 minutes. Discard the supernatant and wash the pellet once with 1×NEB Buffer 3.1. Again, discard the supernatant and resuspend the pellet in 199 μl of 1×NEB Buffer 3.1. Mix well, then add 1 μl of 20% SDS (final concentration 0.1%). Mix well again, place it in a 65°C metal bath for 10 minutes, and then immediately put it on ice for 3 minutes. Finally, add 10 μl of 20% Triton X-100 to neutralize SDS. Mix well by pipetting, and incubate at 37°C with shaking for 15 minutes.

### Enzyme Digestion and ligation

Add 200U of DpnII (NEB, no. R0543L) or 200U of AluI (NEB, no. R0137L) to the reaction solution from the previous step. Mix well and digest overnight at 37°C. Take 2 μl of the solution, add 7 μl of NF water and 1 μl of Proteinase K, then de-crosslink at 65°C for 1 hour. Check the band size by 1% gel electrophoresis.

Wash the sample twice with washing buffer (containing 1× PBS, 1 mM EDTA, 1 mM EgTA, and 0.1% Triton X-100). Resuspend in 195 μl of 1×T4 DNA ligase buffer containing 0.1% BSA. Add 5μl of 400 U/μl T4 DNA ligase (NEB, no. M0202L) to achieve a final concentration of 10 U/μl. Mix well and incubate at 16°C for 4 hours.

Again, take 2 μl of the solution, add 7 μl of NF water and 1 μl of Proteinase K, then de-crosslink at 65°C for 1 hour. Check the band size by 1% gel electrophoresis.

### Enzyme Digestion for single cell barcode attachment

Start by washing the cell nuclei twice with 1.1×Cutsmart buffer, then resuspend in 200 μl of the same buffer. Add 20 μl of HpyCH4V (NEB, no. R0620L) and incubate at 37°C at 1,200 rpm on a shaking mixer for 4 hours.

Take a 2 μl sample, add 7 μl of NF water, and 1 μl of proteinase K. De-crosslink at 65°C for 1 hour and check the band size by 1% gel electrophoresis. After the digestion reaction, centrifuge the cell nuclei at 500g for 3 minutes. Remove the supernatant and wash twice with washing buffer (1× PBS, 1 mM EDTA, 1 mM EgTA and 0.1% Triton X-100).

Resuspend and count the cell nuclei using a cell counting plate. Take 5 × 10^5 cell nuclei into a new 1.5 ml centrifuge tube and add 25 μl of dA-tail reaction buffer and 10 μl of Klenow fragment. Both components are from the NEBNext dA-Tailing Module (NEB, no. E6053L). Make up the volume to 250 μl with NF water, and incubate at 37°C at 1,200 rpm on a shaking mixer for 90 minutes.

Add 200 μl of reaction termination solution (1× PBS, 50 mM EDTA, 50 mM EgTA and 0.1% Triton X-100), then centrifuge at 900g for 2 minutes. Wash twice with washing buffer and resuspend in 200 μl of the same buffer. Filter the cells through 40 μm and 20 μm filters.

Count the cell concentration using a cell counting plate. Take 2 × 10^5 cell nuclei into a new 1.5 ml centrifuge tube. Make up to 1,165 μl with BSA buffer (1× PBS, 0.1% Triton X-100, 0.3% BSA).

### Adding Index Adapter

Use the synthesized 96-strip index primer, designed to avoid the GATC sequence, along with another primer to prepare the 50 μM index-Y adapter. Take 2 μl of each of the 96 adapters and add them to the wells of the 96-well plate containing the cells.

Next, prepare the ligation reagents. Mix 220 μl of 2× Instant Sticky-end Ligase Master Mix (NEB, no. M0370), 352 μl of 5× Quick Ligase Buffer (NEB, no. B6058S), and 132 μl of 1,2-propanediol (Sigma-Aldrich, no. 398039). Distribute the mixture into the wells of the 96-well plate containing the cells and adapters.

Incubate at 20°C at 1,600 rpm on a shaking mixer for 3 hours, shaking for 30 seconds every 5 minutes. After the adapter addition reaction is complete, add 20 μl of reaction termination solution (1× PBS, 50 mM EDTA, 50 mM EgTA and 0.1% Triton X-100) to each well. Shake to mix and let stand at room temperature for 10 minutes to terminate the reaction.

Add an extra 80 μl of reaction termination solution to each well. Combine the products from the 96 wells into a 15 ml centrifuge tube. Centrifuge at 800g for 5 minutes, wash twice with washing buffer, and resuspend in BSA buffer.

Filter the cells through a 10 μm filter to screen out single cells. Count the cell concentration using a cell counting plate. Take 1400 cell nuclei into a new 1.5 ml centrifuge tube, and make up to 350 μl with BSA buffer (1× PBS, 0.1% Triton X-100 and 0.3% BSA). Mix well and distribute into a 96-well plate, with 3 μl per well. The total quantity of cell nuclei in the 96 wells should be about 1,200.

### Reverse crosslinking and PCR Amplification

First, prepare 0.1 U/ml Qiagen protease (QIAGEN, no. 19157) with EB buffer and distribute 3 μl per well. Shake the 96-well plate to mix, then perform flash centrifugation. Place the plate on a PCR machine to perform the decrosslinking reaction (50°C for 1 hour, 70°C for 30 minutes, and 65°C for 2 hours).

Next, add 0.6 μl of 10 μM index-containing i7 PCR primer (5’-CAAGCAGAAGACGGCATACGAGAT - 8bp index - GTCTCGTGGGCTCGG - 3’) to each wel. This primer is designed to avoid the GATC sequence.

Then, prepare the PCR system. This includes 760 μl of 2×KAPA HiFi readymix (KAPA Biosystems, no. KK2602), 65 μl of HiChew-P5 primer (5’-/phos/AATGATACGGCGACCACCGAGATGTACAC - 3’), and 135 μl of NF water. Add 9.3 μl of this mixture to each well.

Afterwards, perform 12 cycles of the amplification reaction on a PCR machine.

Lastly, combine the PCR products from the 96 wells into a 15 ml centrifuge tube. Purify the combined samples using the QIAGEN PCR purification kit (QIAGEN, no. 28104).

### Methylation Marking and IP

Take 1 μg of purified DNA and add it to the reaction consisting of 5 μl 10×dam Methyltransferase Reaction Buffer, 0.25 μl 32 mM SAM, 1ul dam Methyltransferase (NEB, no. M0222L), and water to make up to 50 μl. React at 37°C for 1 hour to perform m6A methylation marking at the GATC site. Then, purify the marked product using 1×Ampure XP beads.

Before DNA IP, Protein A/G Magnetic Beads (Thermo, no. 88802) were washed by taking 10 μl of beads per sample and washing them twice with PBST containing 0.1% tween 20, and resuspend in 50 μl SuperBlock (PBS) Blocking Buffer (Thermo, no. 37580) for 15 minutes at room temperature. Beads were washed twice with 1×IP buffer (10 mM Tris-HCl (pH 7.5), 150 mM NaCl, 0.5% (vol/vol) CA630 and 1 mM EDTA) and resuspend in 48 μl 1×IP buffer. Add 2ul Anti-N6-methyladenosine (m6A) antibody (Merck, no. ABE572-I-100UG) and incubate at 4°C with rotation overnight.

Antibody-coated beads were then washed 5 times with 1×IP buffer and resuspended in 50 μl 1×IP buffer for later use. 500 ng of marked DNA was made up to 40 μl with nuclease-free water, denature at 95°C for 5 minutes, then immediately place on ice for 2 minutes. Add 10 μl of pre-cooled 5×IP buffer (50 mM Tris-HCl (pH 7.5), 750 mM NaCl, 2.5% (vol/vol) CA630 and 5 mM EDTA) and 50 μl of antibody-coated beads. Mix well and incubate at 4°C with rotation for 2 hours.

Place the beads on a magnetic rack to remove the supernatant. Wash once with pre-cooled medium-stringency RIPA buffer (10 mM Tris-HCl (pH 8.0), 300 mM NaCl, 1 mM EDTA, 0.5 mM EGTA, 1% (vol/vol) Triton X-100, 0.2% (vol/vol) SDS and 0.1% (vol/vol) sodium deoxycholate) on ice. Wash twice with pre-cooled high-stringency RIPA buffer (10 mM Tris-HCl (pH 8.0), 350 mM NaCl, 1 mM EDTA, 0.5 mM EGTA, 1% (vol/vol) Triton X-100, 0.23% (vol/vol) SDS and 0.1% (vol/vol) sodium deoxycholate) on ice. Wash once again with pre-cooled medium-stringency RIPA buffer on ice, and finally wash twice with pre-cooled 1× IP buffer on ice.

Add 22 μl of EB buffer to prepare 0.05 U/ml Qiagen protease, and incubate at 50°C for 30 minutes on a shaking mixer. Place the beads on a magnetic rack and transfer 20 μl of the supernatant to a new PCR tube. Incubate at 70°C for 15 minutes on a PCR machine to inactivate Qiagen protease. Add 30 μl of PCR system (25 μl 2×KAPA HiFi readymix, 1.5 μl HiChew-P5 primer, 1.5 μl P7 primer 5’-CAAGCAGAAGACGGCATACGAG-3’, 2 μl water), and perform 6-8 cycles of amplification reaction. Purify the PCR product with 1×Ampure XP beads.

### Library Circulation and Sequencing

Use the MGIEasy Circulating Kit (MGI, no.1000005259) to circularize the library according to the instructions, circularize the product using the MGISEQ-2000RS High Throughput Sequencing Reagent Kit (PE100) (MGI, no.1000012554) to prepare DNA nanoball and sequence on the MGISEQ-2000 platform using the PE100+100+10+10 sequencing mode.

### SCA-seq

SCA-seq were performed as previously description^12^.

### Preprocessing of and bulk Hichew and snHiChew datasets

Raw reads for each cell were sorted using an in-house script. Single-cell or bulk sample Hi-C paired-end reads were then aligned to the hg19 or mm10 reference genome using HiC-Pro (v.3.1.0, default settings). This data was used to generate HiC-Pro filtering and alignment metrics, valid pairs, and contact matrices, which were then corrected with ice_norm. For each chromosome, contact matrices at resolution levels ranging from 10kb to 1Mb were created for further analysis. In order to estimate the number of valid cells in the snHiChew data, a barcode rank plot was used to identify the steep drop-off pattern that separates valid cells from background noise. Specifically, all detected snHiChew barcodes were plotted in descending order based on the number of nonduplicated valid pairs associated with each barcode. The R package kneedle was used to identify the distinction between valid cells and non-cell barcodes. The data associated with the valid cells were then used for further analysis.

### Comparison of bulk chromatin architecture datasets

We summarized key metrics such as the cis/trans ratio, valid pair ratio, valid pair dup, and others from the HiC-Pro statistical outputs. To clarify, the valid pair ratio was computed by dividing the valid pairs (prior to removing duplication) by the reported pair from HiC-Pro. This effectively shows the enrichment efficiency. The valid pair dup is the duplicated segment of valid pairs as identified by HiC-Pro, which evidences the level of sequencing saturation.

To analyze the contact matrix pattern, we first converted the nonduplicated valid pairs from bulk Hichew and other chromatin conformation capturing methods into mcool files using cooler (v0.8.2). We then used the R package HiCExperiment (v1.2.0) ^23^to perform a paired comparison analysis at different contact map resolutions. This analysis offers insights into hi-c features at the whole genome, chromosome, topological associated domains (TAD), and chromatin loop levels.

We measured the correlation of contact matrices at chromosomal and TAD levels using genome-wide eigenvector scores (compartment score) and TAD insulation score, calculated by Cooltools (v0.4.1). We then calculated the corresponding correlation using in-house R scripts and Pearson methods. Lastly, the contact distance decay curve was calculated using HiCExplorer (v3.7.2).

### snHiChew collision rate estimation

The snHiChew HEK293T-NIH3T3 mixture data was processed using a combined genome of hg19 and mm10, using established snHiChew preprocessing techniques. We used nonduplicated valid pairs in the cell rank plot to identify a steep drop-off pattern and pinpoint the empty cell barcodes. For valid cell barcodes, we assigned them to HEK293T if a minimum of 89% of nonduplicated valid pairs were linked to hg19. Similarly, we assigned them to NIH3T3 if at least 89% of nonduplicated valid pairs were associated with mm10. Remaining valid cell barcodes were classified as doublets.

### Clustering of snHiChew data in HEK293T-NIH3T3 mixture data

The corresponding pairs related to the annotated HEK293T and NIH3T3 cell barcodes are combined and marked for grouping. We employed Higashi ^24^ with standard settings for dimensionality reduction calculations. For UMAP clustering, we used the first 15 principal components of Higashi embeddings with parameters set to “n_neighbors=5, min_dist=0.01, metric=’correlation’”.

### Comparison of single cell chromatin architecture datasets

We compared the performance of snHiChew’s chromatin architecture data with published data from Dip-C and snHi-C, well-known single-cell chromatin conformation capture technologies. The datasets were obtained from the National Center for Biotechnology Information (NCBI) Gene Expression Omnibus (GEO) using the codes GSE94489 and GSE146397.

We applied the same preprocessing and HiC-Pro metrics based benchmark procedures to Dip-C, snHi-C, and snHiChew as we previously described. To assess sequencing yield efficiency, we downsampled raw reads to various levels: 25k, 50k, 100k, 200k, 400k, 800k, 1.6M, 3.2M, 6.4M, 12.8M. We only included cells with read numbers that exceeded these thresholds.

We then calculated the number of non-duplicated valid pairs per read for each threshold, fitting the data points to a saturation curve using the model Y = B max * X/(Kd + X). Finally, we estimated the read number for sequencing saturation for each single-cell chromatin conformation capture technology.

The false positive rate was assessed by looking at the unexpected interactions between mitochondrial DNA and nuclear DNA. We computed the false positive rates for each unique valid pair identified by HiC-Pro in every valid cell of the snHiChew and Dip-C datasets, using the method described in Trac-looping ^25^.

### In silico cell phasing over the cell cycle

We carried out cell-cycle analysis following the method detailed in a prior study ^26^. In short, we used the HEK293T 2-phase Repli-seq dataset (4DNESSV33VOL, 4DNESH4XLJCW) obtained from the 4D Nucleome Data Portal to label the early/late repli-score ratio for each cell. A higher early/late repli-score ratio indicates that the cell is nearer to the early S-phase of the cell cycle.

### Three-dimensional genome modeling

We utilized Chrom3D ^27^ to generate three-dimensional structural genome models. The model setup files were created from the HiC-pro raw contact matrix with a 50kb bin. A single cell from each cluster was chosen to symbolize the melting and concretion phases. We adhered to the Chrom3D protocol for the modeling process, which can be found at https://github.com/Chrom3D/pipeline. In the model setup file, three regions of interest (ROI) were distinctly color-labeled. The models were generated with the following settings: Chrom3D -n 100000 -r 5.0 --nucleus -y 0.01 -l 10000 -c 0.001 [Model Setup File] > [Output file (CMM)].

### Statistics

We conducted a two-sided t-test on most of the parametric data, which followed a normal (or log-normal) distribution. For this same data, we also performed a Pearson correlation analysis. We used the Wilcoxon rank test for non-parametric data or data that was not normally distributed.

## Data availability

The data were stored at https://db.cngb.org/search/project/CNP0005635/ and at NCBI BioProject PRJNA1109567. Other public datasets used in this study were downloaded from NCBI GEO with the following accession numbers: ChIP-seq (CTCF ENCSR135CRI H3K4me3 ENCSR000DTU; H3K27ac ENCSR000FCH), snHi-C (GSE94489), Dip-C (GSE146397), HEK293T in situ Hi-C (GSE143465), SCA-seq (PRJNA917827). The source data are provided with this paper.

## Code availability

Custom scripts used in this study are available from https://github.com/genometube/snHiChew.

## Author contributions

CT designed and oversaw the experiments. ZCC conducted the laboratory experiments; YMX conducted the bioinformatics data analysis. All authors collectively performed the data analysis. All authors have read and approved the final draft of the manuscript.

---

**Supplemental Fig1.**
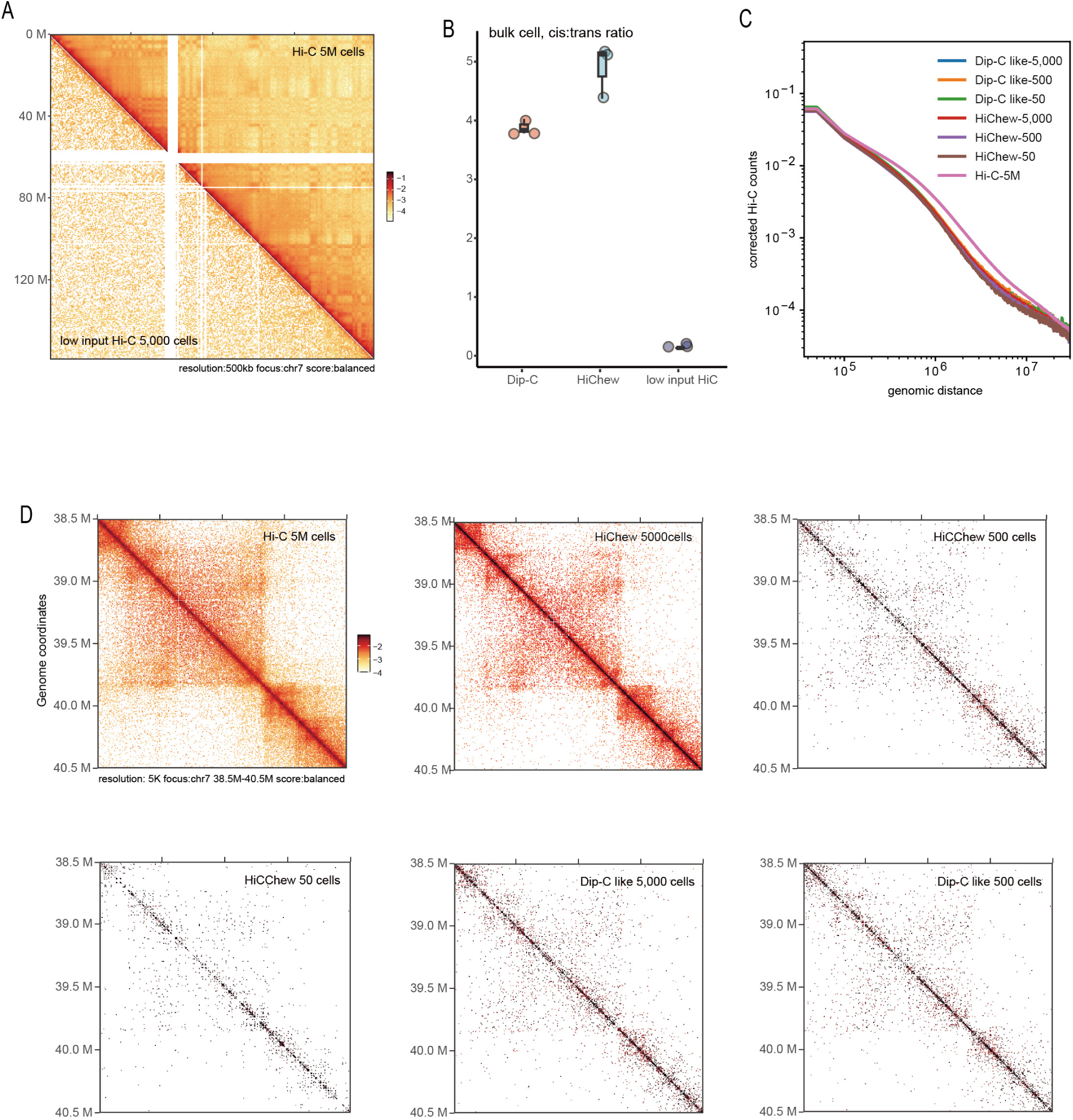
Initial Assessment of snHi-C, HiChew, Dip-C. (A) Contact maps on chr7, comparing Hi-C results from 5000 cells with 5 million cells. (B) Ratio of cis-trans. (C) Interaction distance decay curve. (D) Sample region of loops on chr7, from 38.5M-40.5M, at 5kb resolution.

**Supplemental Fig2.**
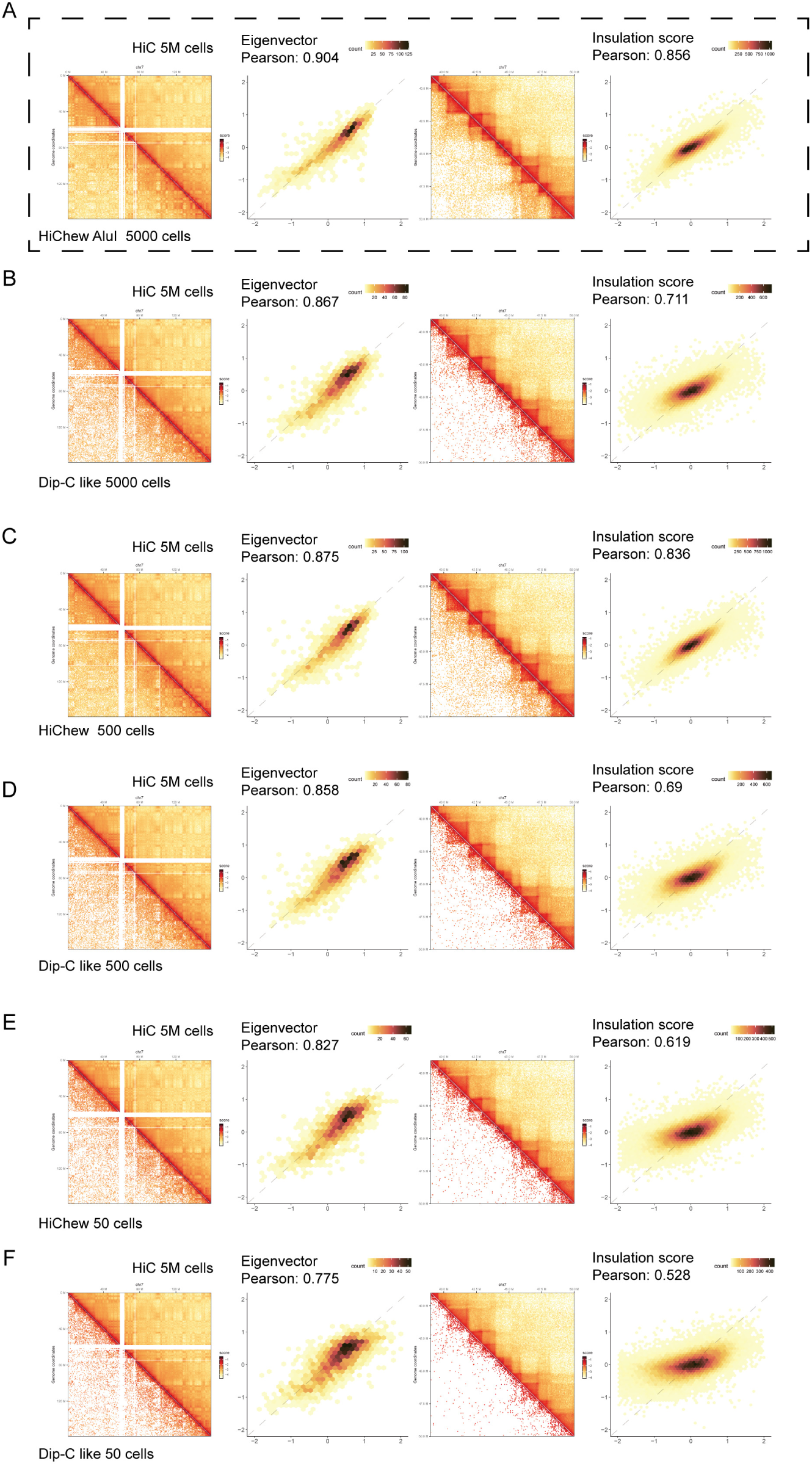
Comparison of HiChew, Dip-C 5000 500 and 50 Cells with established Hi-C 5 million cells standard. (A-C) This presents the HiChew 500 and 50 cells in Chr7 at a 500kb resolution. The correlation of eigenvalues is demonstrated in the lower panel. (B-D) Here we observe the HiChew 500 and 50 cells in Chr7:39M-50M at a 50kb resolution. The lower panel illustrates the correlation of insulation scores.

**Supplemental Fig3.**
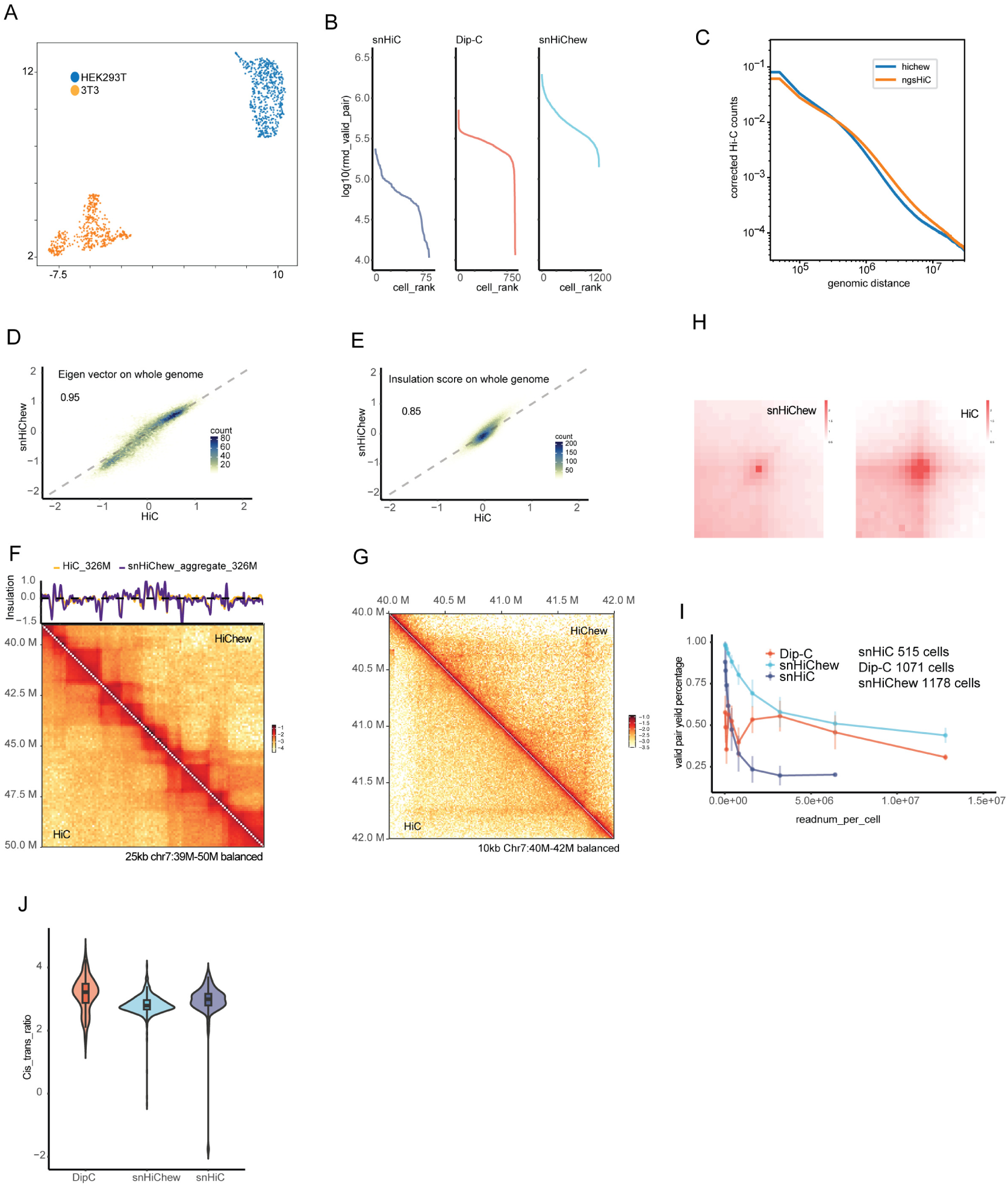
Here is a comprehensive analysis of snHiChew. (A) Higashi software (cite) uses UMAP to embed a mix of 3T3 cells and HEK293T cells. (B) We’ve ranked the valid pairs for each cell from high to low to establish a cutoff for the empty barcodes. (C) This graph shows the distance decay curve of the contact pairs. (D) Here, we show the correlation of the eigenvalue between Hi-C (326M valid pairs) and HiChew (aggregated 1178 cells, 326M valid pairs) on the whole genome. (E) This graph reveals the correlation of the insulation score between Hi-C and HiChew on the whole genome. (F) The top panel displays insulation scores on chr7:39M∼50M, while the bottom panel presents the contact maps at a resolution of 10kb. (G) This is the contact map on chr7:40M-42M. (H) Here is the aggregation plot(APA) of loops. (I) This graph shows the deduped valid pair yield percentages versus the sequencing depths in each cell. (J) The final graph depicts the cis-trans ratio among Dip-C, snHiChew, and snHi-C.

**Supplemental Fig4.**
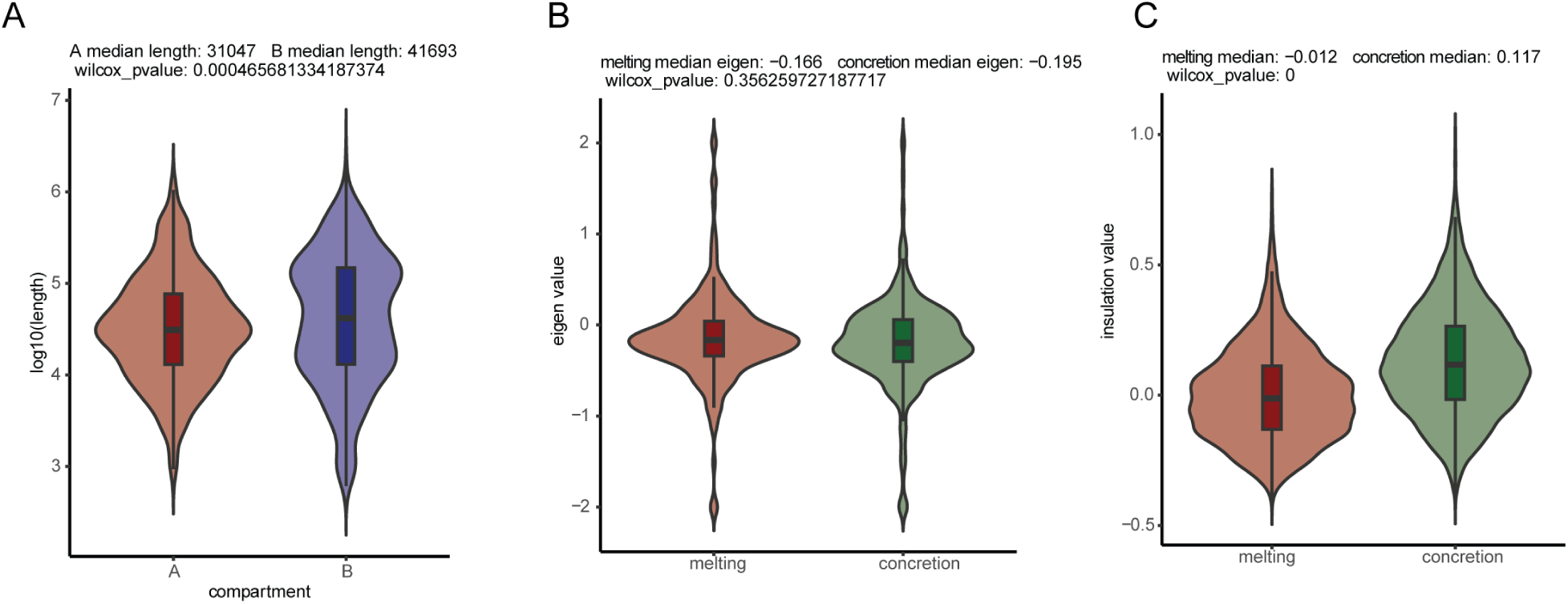
(A) This shows the distribution of coding gene length between the A compartment and the B compartment. (B) Here we have a violin plot of the eigenvalues for the melting and concretion cluster in the area chr7:145MB-148MB. (C) This is a violin plot of the insulation scores for the same melting and concretion cluster in chr7:145MB-148MB.

**Supplemental Fig5.**
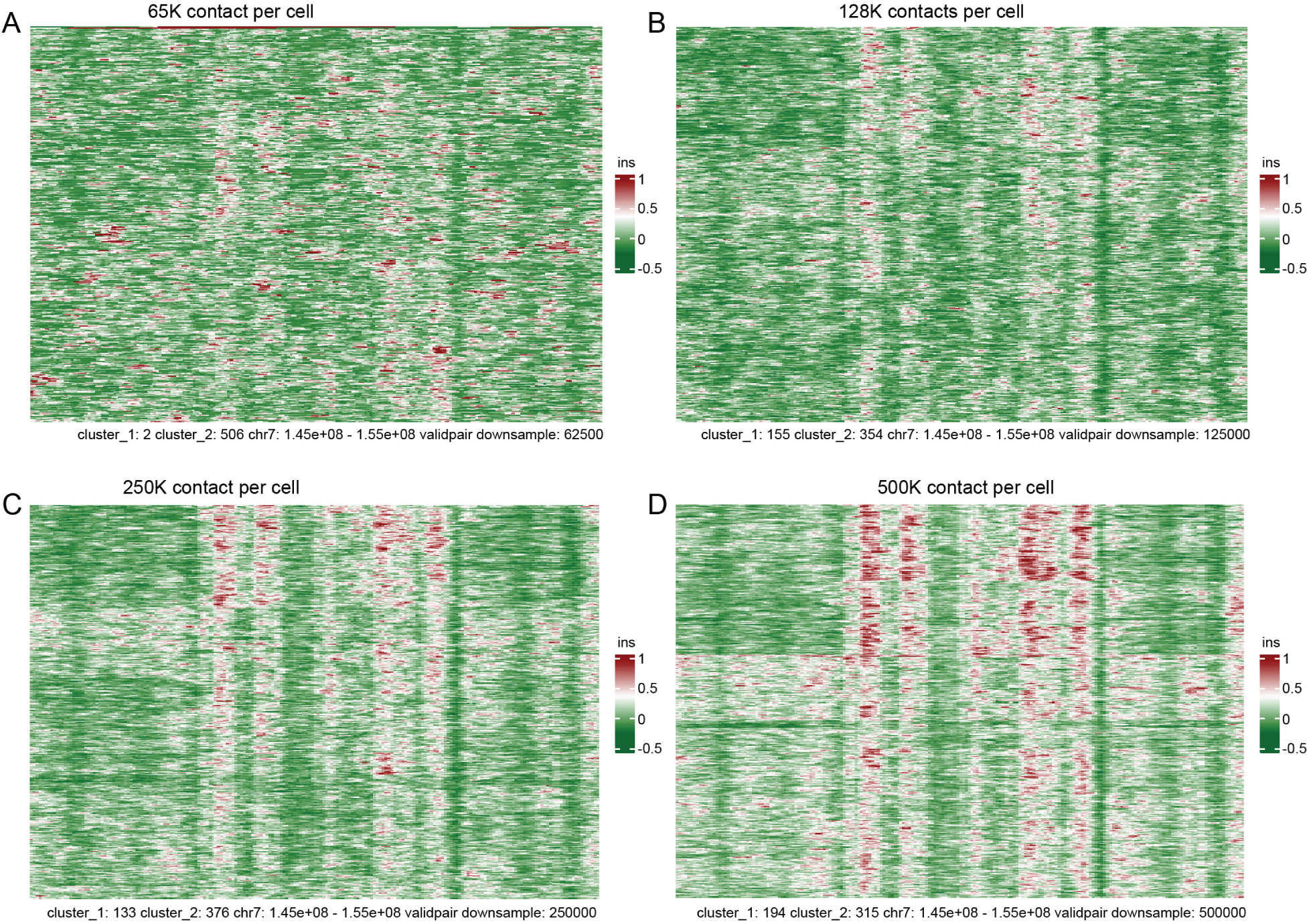
This is a simplified representation (downsampled from 65K to 500K) to better illustrate the effectiveness of cell insulation scores in clustering.

**Supplemental Fig6.**
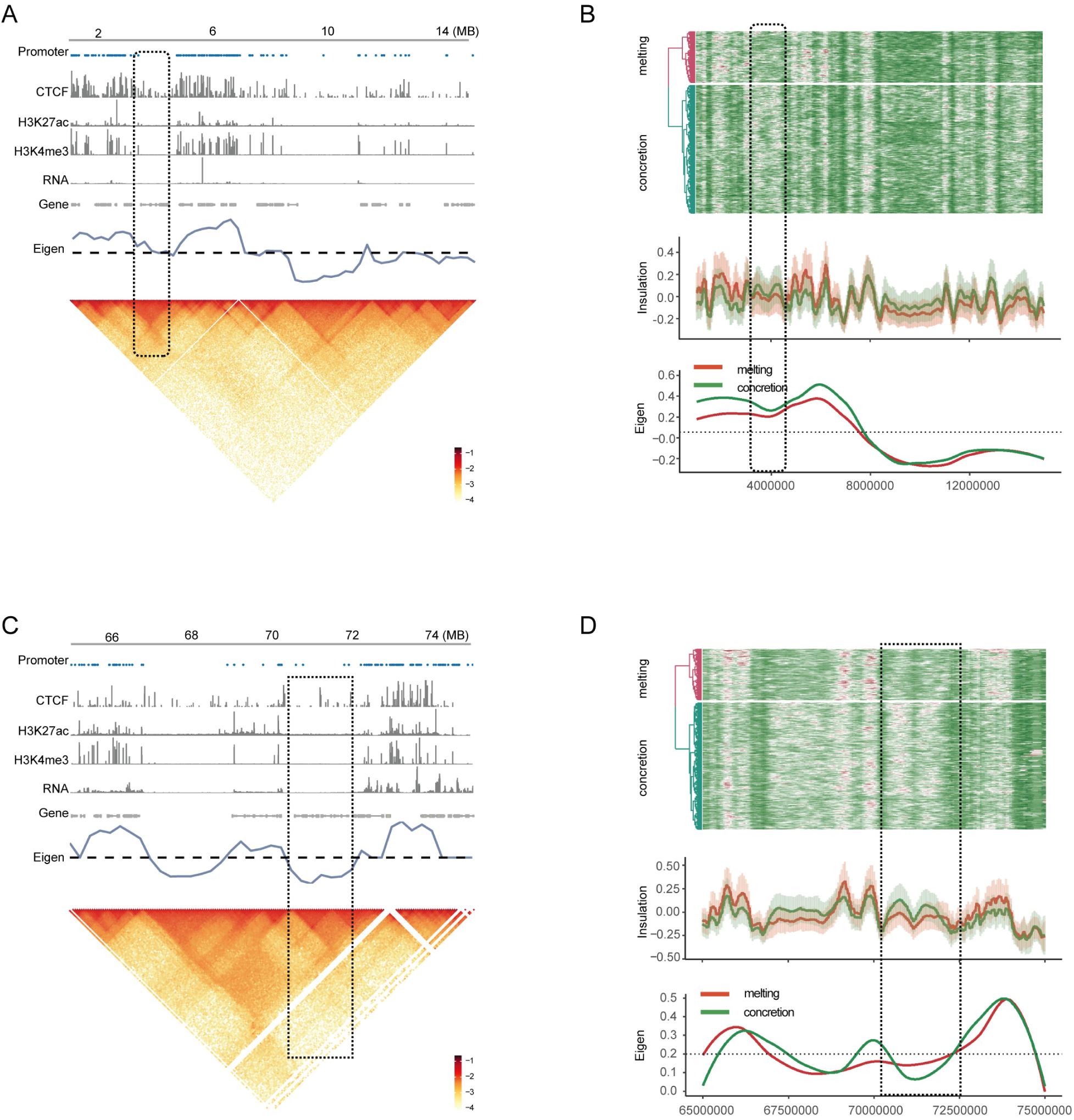
This figure provides data for two other example regions, chr7:1-15MB and chr7:65-75MB. (A,C) We gathered CTCF, H3K27ac, and H3K4me3 peak data from ChIP-seq. We then calculated the eigen value from the contact maps. The term ’promoters’ refers to the location of each gene’s promoter, as shown. The bottom heatmap presents the chromatin contact maps. (B,D) This panel demonstrates the clustering analysis of the insulation score for individual cells. The bottom panel displays the average insulation score and eigen value of the melting and concretion cluster.

**Supplemental Fig7.**
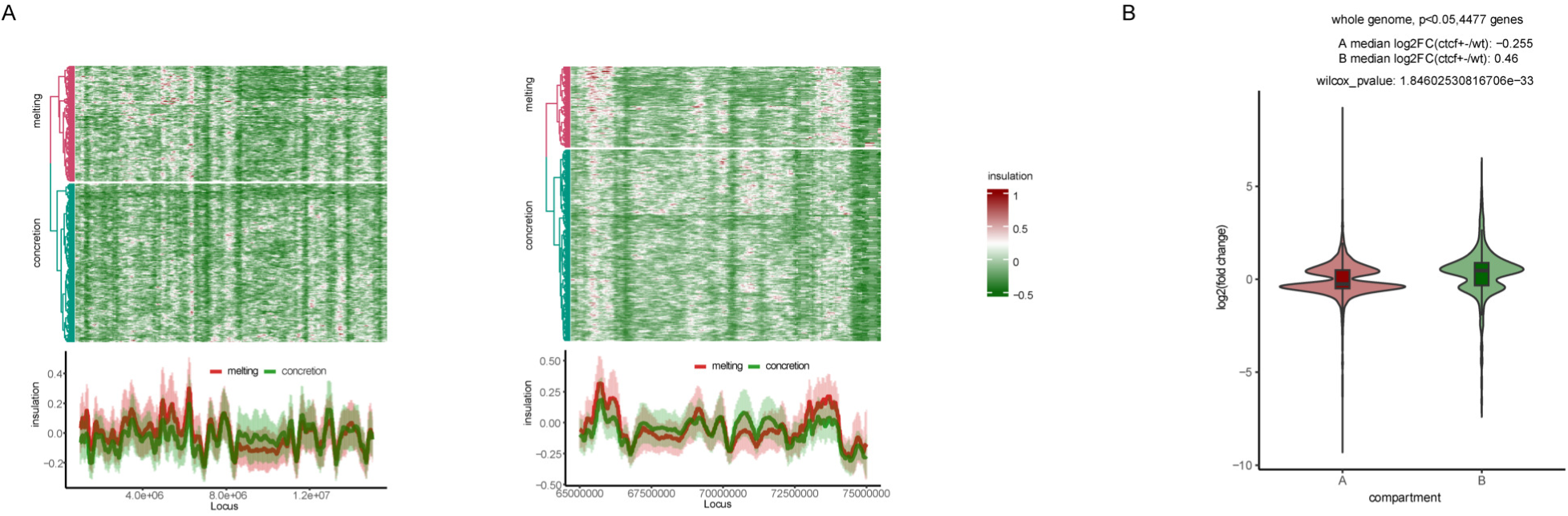
(A) Here are two other example regions in CTCF+/-, chr7:1-15MB and chr7:65-75MB. (B) We’ve classified the significant genes into two categories: B compartment long genes and A compartment short genes. The y-axis represents the log 2 fold changes in gene expression when comparing CTCF+/- to WT.

**Supplemental Fig8.**
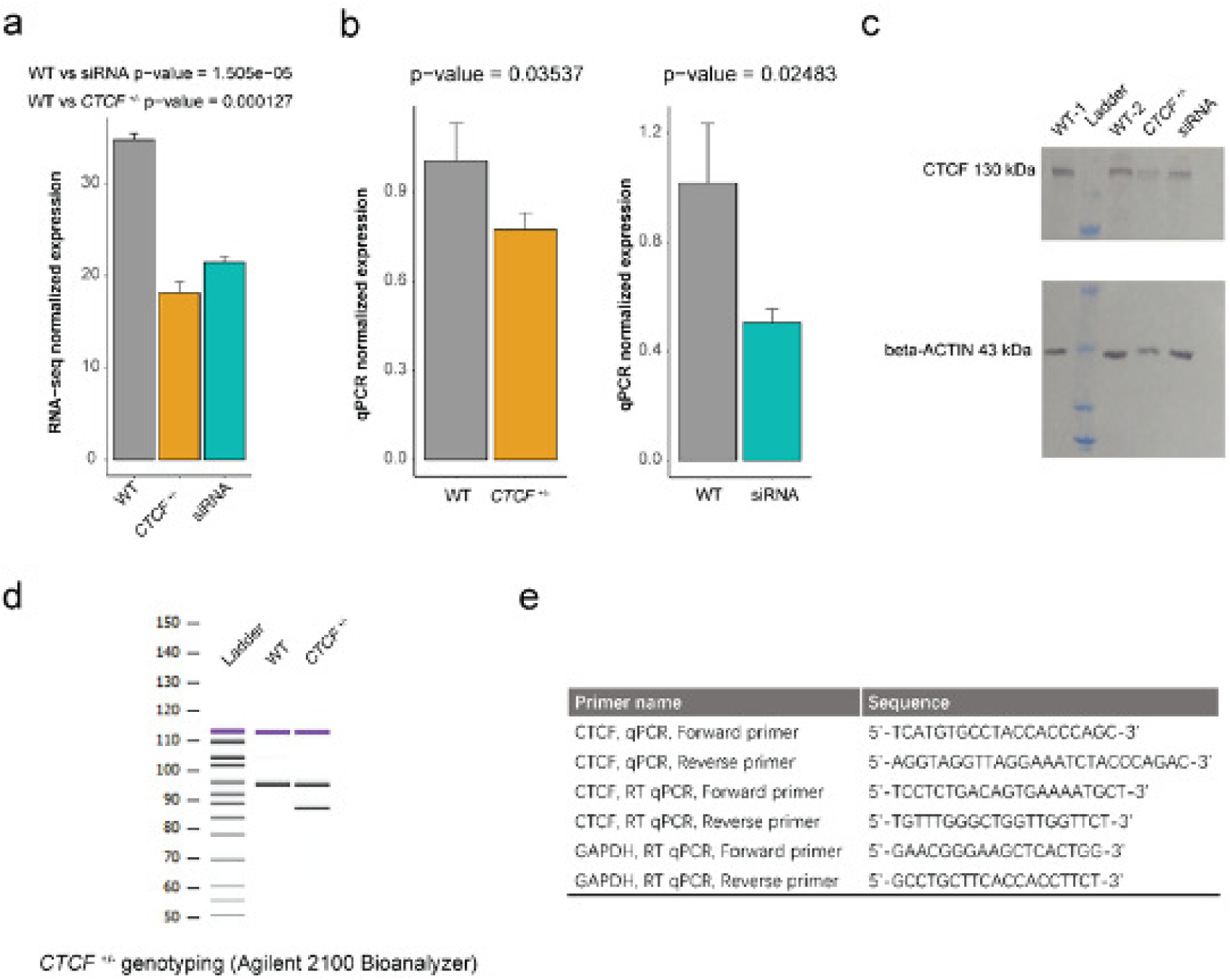
Cell phenotype characterization. (A) The counts of *CTCF* gene in RNA-seq. (B) The qPCR normalized by *beta-ACTIN*. (C) The western plot of CTCF normalized by *beta-ACTIN*. (D) The genotyping of the cell lines. (E) The primers of genotyping.

